# A novel in-vivo phagocytosis assay to gain cellular insights on sponge-microbe interactions

**DOI:** 10.1101/2023.02.28.530395

**Authors:** Angela M. Marulanda-Gomez, K Kristina Bayer, Lucia Pita, Ute Hentschel

**Author notes:** **Correspondence:** Lucia Pita, Ute Hentschel.

## Abstract

Sponges harbor diverse, specific, and stable microbial communities, but at the same time, they efficiently feed on microbes from the surrounding water column. This filter-feeding lifestyle poses the need to distinguish between three categories of bacteria: food to digest, symbionts to incorporate, and pathogens to eliminate. How sponges discriminate between these categories is still largely unknown. Phagocytosis is conceivable as the cellular mechanism taking part in such discrimination, but experimental evidence is missing. We developed a quantitative in-vivo phagocytosis assay using an emerging experimental model, the sponge *Halichondria panicea*. We incubated whole sponge individuals with different particles, recovered the sponge (host) cells, and tracked the particles into the sponge cells to quantify the sponge’s phagocytic activity. Fluorescence-activated cell sorting (FACS) and fluorescent microscopy were used to quantify and verify phagocytic activity (i.e., the population of sponge cells with internalized particles). Sponges were incubated with a green microalgae to test the effect of particle concentration on the percentage of phagocytic activity, and to determine the timing where the maximum of phagocytic cells are captured in a pulse-chase experiment. Lastly, we investigated the application of our phagocytic assay with other particle types (i.e., bacteria and fluorescent beads). The percentage of phagocytic cells that had incorporated algae, bacteria, and beads ranged between 5 to 24 %. We observed that the population of phagocytic sponge cell exhibited different morphologies and sizes depending on the type of particle presented to the sponge. Phagocytosis was positively related to algal concentration suggesting that sponge cells adjust their phagocytic activity depending on the number of particles they encounter. Our results further revealed that sponge phagocytosis initiates within minutes after exposure to the particles. Fluorescent and TEM microscopy rectified algal internalization and potential digestion in sponge cells, and suggests translocation between choanocyte and archeocyte-like cells over time. To our knowledge, this is the first quantitative in-vivo phagocytosis assay established in sponges that could be used to further explore phagocytosis as a cellular mechanism for sponges to differentiate between different microorganisms.

## Introduction

Early branching metazoans provide an opportunity to investigate the evolution of host-microbe interactions. Sponges (phylum Porifera) are benthic suspension feeders that actively filter large volumes of water. Due to this filter-feeding lifestyle, they are constantly exposed to high numbers of particles, which poses the question of how sponges distinguish and process between the different types of microbes they encounter. Sponges exhibit three well-defined types of epithelial layers: the external pinacoderm composed of T-shaped or flatted cells (i.e., pinacocytes) which covers the outside of the animal; the internal choanoderm containing the flagellated cells (i.e., choanocytes); and the mesohyl, a collagen-like matrix within which other sponge cells, skeletal components, and most symbiotic microbes reside (1,2). These filter feeders evolved a complex branched water canal system (i.e., the aquiferous system) comprised of several choanocytes arranged into hollow chambers which generate water flow by the beating of their flagella. Water enters the sponge through small pores, or ostia, which spread along the animals’ outer surface, circulates through the incurrent canals into the choanocyte chambers, and exits through larger outflow openings, or oscula. The particles (e.g., bacterio- and phytoplankton) that are filtered by the choanocytes are translocated to amoebocyte-like cells (i.e., archeocytes) which are regarded as potential nutrient transporters along the sponge (3–5). The participation of both choanocytes and archeocytes in intracellular digestion is supported by single-cell transcriptomic data which shows enrichment in genes related to e.g., engulfment and motility, lysosomal enzymes, and phagocytic vesicles (6,7). Despite this filter-feeding lifestyle, sponges harbor a highly diverse community of associated microbes in their mesohyl that is remarkably different from the bacterioplankton in the surrounding seawater (8–10). It remains enigmatic if, and how, sponges distinguish between food to digest, symbionts to acquire, and pathogens to eliminate.

Selectivity in particle uptake continues to be a controversial topic in sponge physiology. Some studies regard particle size as the main parameter driving the selection of microbes during the filtering process (11–14), whereas others propose a size-independent discrimination between particles involving individual particle recognition, sorting, and transport through the sponge tissue during the digestion process (5,12,15–17). Furthermore, two feeding experiments with culturable bacteria provide first evidence that sponges take up seawater bacteria but not sponge-associated bacteria (18,19). The latter studies hypothesized that the lower uptake rates of sponge-associated bacteria isolates result from slime capsules or secondary metabolites that protected symbionts from being recognized and ingested by the sponge. How the sponge host exactly discriminates microbes at the cellular level is still unclear, yet we posit that this cellular host-microbe recognition mechanism must be essential for establishing and maintaining symbiotic interactions, and a stable microbiome in sponges.

We thus hypothesize that phagocytosis will play a major role in sponge-microbe interactions. Hijacking of phagocytosis promotes the colonization and maintenance of microbes during symbiotic interactions (20). Phagocytosis includes the recognition and ingestion of particles larger than 0.5 µm within a plasma-membrane envelope (i.e., phagosome) (21). It is a highly conserved cellular process from unicellular to multicellular organisms, involved in nutrition, defense, homeostasis and symbiosis (20,22,23). Symbionts from diverse hosts such as amoeba, leech and squid are capable of escaping different stages of the phagocytic process, avoiding either incorporation or digestion by host cells (e.g., 24–26), in their way to colonize and persist in the animal host.

Sponge-associated bacteria are enriched in ankyrin proteins compared to non-associated sponge bacteria (27–30). These proteins modulate host phagocytosis in humans assisting the infection and intracellular survival of symbionts and pathogens (31–33). Experimental evidence on how ankyrins modulate phagocytosis is only probed in non-sponge systems, due to the lack of functional assays in sponges. Free-living amoeba and murine macrophages share physiological and structural similarities to phagocytic sponge cells (i.e., amoebocytes) and have therefore been used as models to study the role of ankyrins in sponges. Interestingly, bacteria expressing ankyrin-repeat proteins were phagocytized, but not digested by the amoeba *Acanthamoeba castellanii* (25), whereas bacteria decorated with phage encoded ankyrins evaded phagocytosis in murine macrophages (24). These pioneering studies serve as evidence that the host phagocytic machinery is targeted by microbes during the symbiotic process and highlights the importance to develop an assay that will allow testing these functions in the sponge host.

Feeding experiments with different types of particles shed some light on the filtration, ingestion, and digestion of particles by sponges (4,5,16). However, the laborious and time-consuming microscopic observations in combination with the lack of well-established methodologies to study cellular processes in sponges hampers our current understanding of the implications of phagocytosis on sponge-microbe interactions. In the closely related phyla of Amoebozoa and Cnidaria advances in genetic manipulation, isolation of symbiont strains and identification of cellular markers facilitated the validation of phagocytosis as the underline mechanism for microbe discrimination (e.g., *Dictyostellium* sp., *Aptasia* sp., *Pocillopora damicornis*, and *Nematoestella vectensis*) (34–39). In hexacorallians, fluorescence-activated cell sorting (FACS) and microscopy allowed further identification and mechanistic characterization of specialized phagocytic cells capable of lysosomal degradation and production of reactive oxygen species (ROS) upon microbe engulfment (38). The development of methods and establishment of model systems are thus fundamental for studying the function of host phagocytosis in microbe selectivity.

In the present study, we describe the development of an in-vivo phagocytosis assay using the Baltic Sea sponge *Halichondria panicea*. We combined live sponge incubation experiments followed by sponge cell dissociation to quantify the uptake of different particles into *H. panicea* cells. Fluorescence-activated cell sorting (FACS), and fluorescence and transmission electron microscopy was performed to gain mechanistic insights into sponge cell phagocytosis. To our knowledge, this is the first quantitative in-vivo phagocytosis assay established in sponges which could be used to further explore the mechanistic underpinnings of recognition and differentiation between sponges and diverse microorganisms, whether food, friend, or foe.

## Material and Methods

### Sponge collection

Individuals of the breadcrumb sponge *Halichondria panicea* (Pallas, 1766) were collected at the coast of Schilksee (54.424278 N, 10.175794 E; Kiel, Germany) in Aug 2022. Sponges were collected at water depths of between 1-3 m and were carefully removed from the crevices of a breakwater structure using a metal spatula. The individuals were directly transferred to the KIMMOCC climate chamber facilities at GEOMAR Helmholtz Centre for Ocean Research (Kiel, Germany), and maintained in a semi-flow through aquaria system supplied with sand-filtered seawater pumped from the Kiel fjord at 6 m depth. The sponges were cut with a scalpel into approximately equal-sized fragments (3.9 ± 1.0 g wet weight [average ± S.D.]), cleared of epibionts, and randomly placed in 10 L aquaria on top of clay tiles (6.3 × 1 cm - Ø x H) for sponges to attach (3-4 sponge fragments per tank). The water flow in the tanks was 0.5 L min^-1^ and a mini-pump with a maximum flow rate of 300 L h^-1^ (Dupla TurboMini) enhanced further water movement in each tank. Sponges were left to heal and attach for 8 days at 10°C room temperature, 17°C water temperature, and a salinity of 16 PSU.

### Tracer preparation for the in-vivo phagocytosis assay

The particles included the microalgae *Nannochloropsis* sp., the *Vibrio* sp. isolate PP-XX7 (16S rRNA gene sequence similarity of 98.6 % with *Vibrio* sp. NBRC 101805 and 97.0% to *Vibrio variabilis* R-40492T), and 1 µm carboxylated beads. We chose these particles because of their size (1-3 µm), clear fluorescence signal, and ecological relevance (i.e., the *Vibrio* strain was isolated from our sponge collection site). The *Nannochloropsis sp*. live culture was purchased from BlueBio Tech (Germany). The stock algal culture concentration was 12 × 10^9^ algae cells mL^-1^, and was kept in the fridge, protected from light until the experiments were performed, as recommended by the manufacturer. The bacteria culture was prepared by inoculating 100 mL of liquid marine broth (Zobell 2216) with a culture that grew for 24h in agar plate. The liquid culture was incubated in the shaker at 120 rpm, at 25°C, for 48 hours until the culture reached the mid to late exponential phase. After this incubation time, the culture OD_600_ was measured (OD_600_ = 1.45) to estimate the aimed bacteria concentration of approx. 10^5^-10^6^ bacteria mL^-1^. The culture was centrifuged at 5000 x g for 5 min to recover the bacteria pellet, resuspended in 0.22 μm filtered artificial seawater, and stained the same day of the experiment with the fluorescent dye 5(6)-TAMRA/SE™ (Thermo Fisher Scientific, C1171) at a final concentration of 1 μM (as in Wehrl et al. 2007). The bacteria suspension was incubated with the dye for 90 min in the dark at room temperature, then centrifuged at 5000 x g for 5 min to wash-off the excess of dye, and the bacteria pellet was resuspended in 0.22 μm FASW. The bacteria suspension was kept at 4°C in the dark until the experiment was performed. The bead stock solution (Fluoresbrite YG microsphere, Cat. 17154-10, Polyscience) with a concentration of 4.55 × 10^10^ particles mL^-1^, was sonicated for 5 min and vortexed immediately before the experiment.

### In-vivo sponge phagocytosis assay

The phagocytosis assay consisted of incubating individual fragments of *H. panicea* with a specific particle for 30 min and tracking its incorporation into the sponge (host) cells. The sponges attached to the ceramic tiles were placed in individual 1 L straight-sided polypropylene wide-mouth Nalgene bottles (ThermoFisher; cat. no. 2118-0032) filled with natural seawater. The tiles laid on a PVC support under which a magnetic stirring was positioned, and the bottles were placed on top of a stirring plate to ensure constant water flow and uniform mixing of the particle during the incubation. Incubations were performed in a climate chamber in which temperature was kept at 10°C. In all cases, sponges incubated in the absence of tracer particles and natural seawater incubations served as controls (n = 4 biological replicates per control and sponges).

First, we tested which is the optimal particle concentration to use during the assay to detect phagocytic cells, we incubated the sponge with three different *Nannochloropsis sp*. concentrations. We added 10 µL, 100 µL, or 1000 µL of the live algal stock culture to each incubation chamber to get a final concentration of approx. ×10^5^, ×10^6^ and ×10^7^ algae mL^-1^, respectively. Secondly, to obtain first insights into the phagocytic process of *H. panicea* we performed a pulse-chase experiment using *Nannochloropsis* sp. to see when we could capture the highest percentage of phagocytic cells. Sponges were incubated independently with algae (10^6^ algae cells mL^-1^ final concentration) over a 30 min pulse phase, and then transferred to the flow-through aquaria with clean natural seawater for a 30 min and 150 min chase phase. Sponges were sampled at t = 30 min (immediately after the pulse phase was over), at t = 60 min and at t = 180 during the chase phase (n = 4 biological replicates per time point). Lastly, we aimed to extend the application of our phagocytic assay to other particles. We incubated independently *H. panicea* individuals with TAMRA-stained bacteria and fluorescent bead solution (approx. final start concentration 10^5^ bacteria cells mL^-1^ and 10^6^ beads mL^-1^, respectively. n = 4 biological replicates per tracer).

### Particle uptake estimation

We expected that particle uptake would influence our quantification of phagocytic sponge cells. Therefore, we assessed whether the sponges were taken up the particles via filtration from the water column during each in-vivo assay. Water samples (1.8 mL) were taken along the incubation period at 0 min, 2 min, 5 min, 7 min, 14 min, 22 min and 30 min. The sampled water was immediately fixed with a mixture of paraformaldehyde and glutaraldehyde in 1x PBS (final concentration 1% and 0.05%, respectively), kept in the dark for 30 min, and stored at -80°C until flow cytometry analysis was performed (following (40)). Algal, bacterial, and bead concentrations from the water samples were estimated using the Accuri C6 Plus flow cytometer (BD Biosciences). Each sample was run for 1 min at a flow rate of 14 µL min^-1^. The cell populations of interest were identified based on the fluorescence of each tracer particle using the BD Accuri C6 Plus Software. The regression fits used for estimating particle uptake can be found in (Pangea PDI-34177 doi xxxx).

### Sponge cell dissociation, preparation, and staining

Immediately after each in-vivo assay, the *H. panicea* fragments were used to extract the sponge cell fraction by tissue dissociation and centrifugation methods (adapted from (19,41)). All buffers and solutions used during the dissociation were adjusted to the salinity and pH of the aquaria at the moment of running the experiment to prevent the cells undergo an osmotic or pH shock. The entire sponge fragments were rinsed in sterile, ice-cold calcium- and magnesium-free artificial seawater (CMFASW), and subsequently cut with a disposable scalpel into small pieces while removing any leftover epibionts or dirt. The tissue fragments were transferred into 50 mL sterile Falcon tubes prefilled with 25 mL of sterile fresh ice-cold CMFASW containing EDTA (25mM) and incubated on ice while gently shaking the tubes horizontally for 15 min. Samples were filtered through 40 μm cell strainers (Corning Inc.) into 50 mL sterile Falcon tubes by gently squeezing the tissue against the walls of the sieve with sterile forceps to remove dissociated tissue fragments and spicules. Ice-cold CMFASW was added to the resulting cell suspension until reaching a total volume of 25 mL and the samples were centrifuged for 5 minutes at 500 x g at 4°C in a swinging router. The supernatant was discarded, and the sponge cell pellet was resuspended in 4 mL of fresh sterile ice-cold CMFASW resulting in a total sponge cell suspension of approx. 5 mL, which was divided into 1 mL aliquots in sterile Eppendorf tubes. The aliquots were fixed with PFA in CMFASW (final concentration 4%) at 4°C in the dark overnight. The fixative was washed off from the cells by centrifugation for 5 minutes at 500 x g at 4°C. Finally, the pellet was resuspended in 1 mL of ice-cold CMFASW. The concentration of sponge cells for each sample was estimated by an automated cell counter (Fluidlab R-300, Anvajo) and adjusted to approx. 5×10^7^ cells mL^-1^ by diluting the cells with ice-cold CMFASW. The fixed cell suspensions were stored at 4°C in the dark for later FACS quantification. For each sample, 200 µL of cell suspension and 4 mL of ice-cold CMFASW were filtered through a 40 μm cell strainer into a 5 mL round-bottom Falcon tube. From this filtered cell suspension, 2 mL were set aside as a control sample into a new tube (i.e., non-stained aliquot). The remaining 2 mL cell suspension was stained with 14 µL of 100ng mL^-1^ DAPI staining (final concentration 0.7 ng uL^-1^). The cells were gently mixed by pipetting and incubated for 30 min at 4°C in the dark.

### FACS quantification of phagocytic active cells

We define phagocytic active cells as those that had incorporated the tracer particle presented to the sponge during the incubation. Sponge cells were analyzed on a MoFlo Astrios EQ^®^ cell sorter (Beckman Coulter) fitted with a 70 µm nozzle and 355nm, 488nm, 561nm and 640 nm wavelength lasers to identify and quantify phagocytic cells. All samples were run three times (technical replicates), for 1 min, and the voltage and pressure were adjusted to ensure to record a similar number of events per second. The gating was performed with the Summit software (V6.3.1) based on the emission of DAPI stained aliquots and the fluorescence of the respective particle used during the phagocytic assay. The strategy we developed for detecting and quantifying the phagocytic active sponge cells of interest followed the subsequent steps. First, to differentiate debris from the “bulk” sponge cells we compared the DAPI emission of non-stained cell aliquots with the DAPI stained aliquots by using the 355 nm UV laser and a filter with a band pass of 448/59 nm (Fig. 1A). A scatter plot was created by selecting the DAPI filter channel on the x-axis and the forward scatter (FCS) on the y-axis, and the DAPI positive cell population was gated. Second, to identify the relative percent of “bulk” sponge cells that had incorporated *Nannochloropsis* sp. cells, the DAPI gate was used to create a new scatter using the laser (561 nm) and filter settings (692/75 nm) in the y-axis to detected algae fluorescence (Fig. 1B). This approach allowed us to distinguish between sponge cells that had incorporated algae from the rest of the sponge cells (i.e., phagocytic from non-phagocytic sponge cells, respectively) by comparing the fluorescence of the control sponges (i.e., without algae) with the sponges provided with algae (Fig. 1B). To identify sponge cells that had incorporated TAMRA-labelled *Vibrio* sp. bacteria from the other sponge cells a scatter plot was created using the side scatter (SSC) on the x-axis and the bacteria fluorescence (laser 561 nm and filter 692/75 nm) in the y-axis (Fig. 1C). Whereas sponge cells phagocytizing beads were detected by plotting the particle size [forward scatter (FCS): x-axis] against the bead fluorescence (laser 488 nm and filter 526/52 nm: y-axis) (Fig. 1D). In all cases, the phagocytic and non-phagocytic sponge cell populations were gated and sorted directly onto microscopy slides (2k-3k cells) for microscopy inspection thanks to the fluorescence of each tracer (Fig. 1). After this verification, the number of events of each gate was used to estimate the relative (%) phagocytic and non-phagocytic cell fraction in relation to the total number of events from these two gates (the complete data set can be found in Pangea PDI-34177 doi xxxx).

**Fig. 1.**
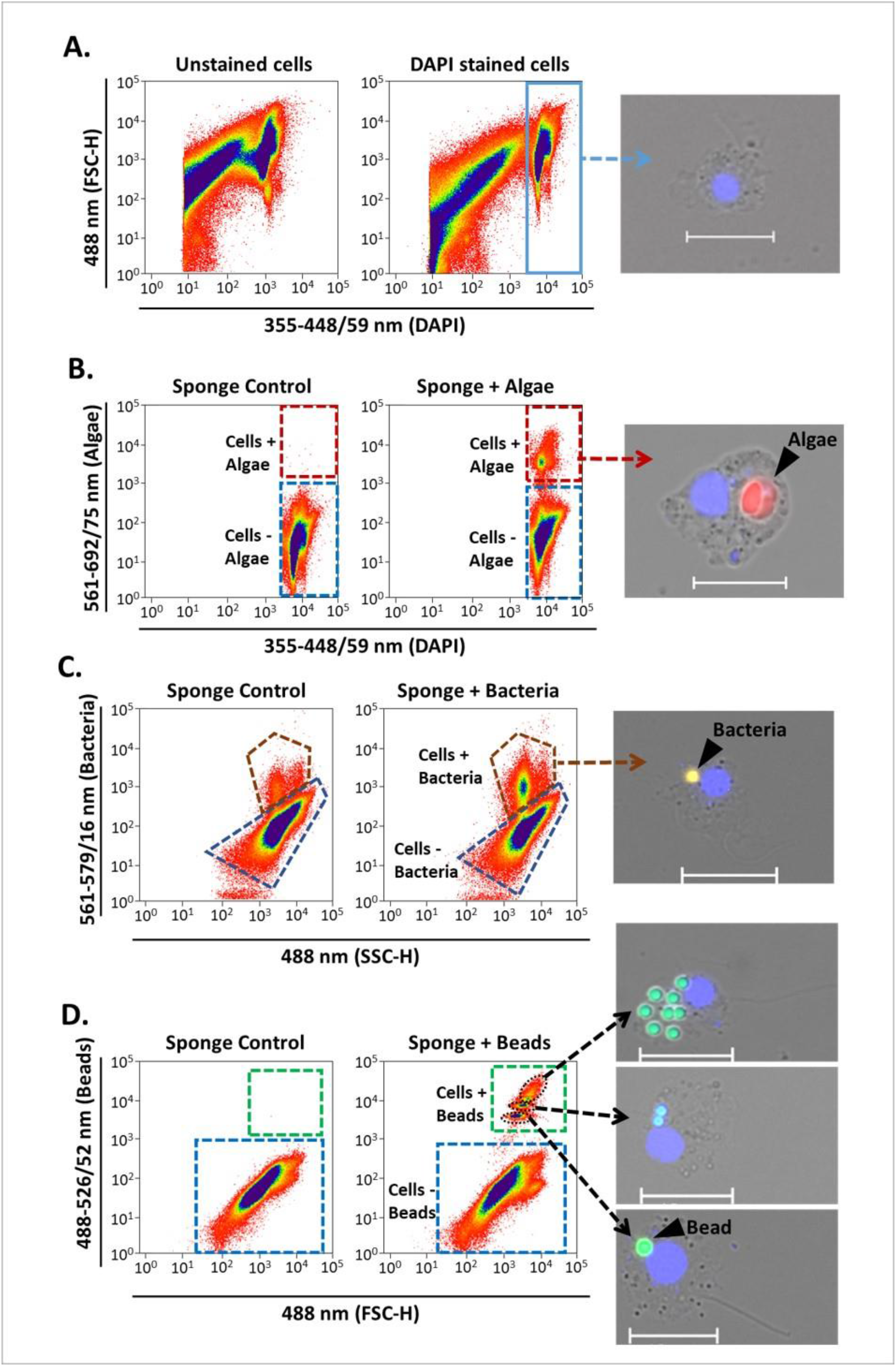
Sponge cell gating strategy for identification and quantification of phagocytic active sponge cells using FACS. Representative cytograms show **(A)** identification of the “bulk sponge cells” population after DAPI staining (blue rectangle). Gates of phagocytic cells with (+) incorporation of **(B)** algae, **(C)** bacteria, and **(D)** beads (red, brown, and green dashed outlines, respectively). Fluorescent microscopy pictures of sorted cells to verify the different gates. Sponge cell nuclei (blue) stained with DAPI. Scale bars: 10 µm. Gates for cells without (-) particle incorporation are shown in each case (blue dashed outlines). Sponges incubated without particles served as controls.

### Fluorescence microscopy

A sub-sample of the sponge cell suspension were inspected by fluorescent microscopy to gain insights into the types of cells involved in the phagocytic process. The cell suspension was mounted in microscopy slides using ROTI mount FluorCare DAPI and 20 × 20 mm cover slides. The slides were air-dried for 30 minutes and examined under an inverted fluorescence microscope equipped with a camera (Axio Observer Z1 with Axiocam 506 and HXP-120 light; Zeiss), at a total magnification of 100x. The following filters and excitations were used: 49 DAPI (335-383 nm for sponge nuclei), 00 Propidium Iodide (530-585 nm for *Nannochloropsis sp*.), 43 HE DsRed (538-562 nm for TAMRA-stained *Vibrio sp*.), and 38 HE GFP (450-490 nm for fluorescent beads). Pictures were acquired using the ZEN Blue Edition software (Zeiss).

### Transmission electron microscopy

In order to locate the particles and verify algal internalization into the sponge cells, a small portion of tissue from two individuals per treatment was collected. The tissue was fixed for transmission electron microscopy (TEM; in 2.5% glutaraldehyde in 1x PBS) overnight at 4°C. Samples were washed three times with 1x PBS for 15 min and then postfixed with 2% osmium tetroxide for 2 h at room temperature, gently shaking the tissue. The sponge tissue was rinsed three times on ice for 15 min and partial dehydration was performed with an ascending ethanol series (2x 30%, 2x 50%, and 2x 70%). Sponge pieces were treated with 5% hydrofluoric acid (in 70% ethanol) overnight at room temperature to remove any silica spicules from their skeleton. Subsequently, samples were rinsed with ethanol (x6 70%), further dehydrated through an increasing series of ethanol, and after embedded in Epon epoxy resin. Ultrathin sections were obtained from region of interest using a Ultracut UC7 (Leica Microsystems) and TEM analysis were performed using a J1010 (Jeol) on the TEM-SEM Electron Microscopy Unit from Scientific and Technological Centers (CCiTUB), Universitat de Barcelona.

### Statistical analyses

Differences in particle uptake between sponge and control incubations were analyzed using an exponential regression approach (as in (42–44)). In each case, the concentrations were corrected based on the average initial concentration of all incubations. The standardized data was fitted to an exponential model (as proposed by (45)) and this was used to estimate the concentration of particles at the start and end of the incubation. To test the effect of algal concentration, chase-time and particle type on the percentage of sponge phagocytic cells, One-way ANCOVA analyses were performed. Particle uptake was used as covariate as we expected that this would influence the estimation of phagocytic sponge cells. ANCOVA assumptions were checked, and significance was determined at the α = 0.05 level. Statistical analysis was performed in R-studio (V4.2.1; Rstudio Team 2022) by fitting an analysis of variance model (aov () function).

## Results

We estimated the sponge’s phagocytic activity as the percentage of sponge cells with internalized fluorescent particles over sponge cells without by coupling incubation experiments with whole *H. panicea* individuals with sponge cell dissociation and FACS. Sponges were incubated with three types of fluorescent particles: *Nannochloropsis* sp., TAMRA-stained *Vibrio* sp., and fluorescent latex beads.

Our optimized cell dissociation protocol for *H. panicea* host cells yielded an average recovery of 1.4×10^7^ ± 5.5×10^6^ cells g^-1^ (sponge wet weight) (Fig. S1). The recovered cell suspension was analyzed by FACS (Fig 1). DAPI signal (stained host nuclei) allowed to identify sponge cells (X-axis), and phagocytic active vs non-phagocytic active cells were distinguished by the fluorescence of the respective particle used during the phagocytic assay (Y-axis). In this way, we quantified phagocytosis as the relative number (%) of phagocytic active (+fluorescent signal) to the total of DAPI+ cells. Phagocytic activity was verified by cell sorting the gated sponge cell populations and fluorescence microscopy only at the beginning of each analysis (Fig. 1).

### Testing tracer concentration for the phagocytic assay

We tested the effect of different tracer concentration on phagocytosis and particle uptake. The total amount of *Nannochloropsis* sp. cells (mean ± SD throughout the text, unless stated otherwise) taken up by *H. panicea* was 4.9 × 10^4^ ± 2.5 × 10^4^ cells and 2.4 × 10^5^ ± 3.5 × 10^5^ cells (Fig. 2A) during the incubation with low (× 10^5^ cells mL^-1^) and medium (× 10^6^ cells mL^-1^) algae concentration, respectively. This uptake corresponds to a reduction in algal concentration of 19.6 ± 9.4 % and 12.2 % ± 14.6 %, respectively. The algal uptake for the highest (× 10^7^ cells mL^-1^) *Nannochloropsis* sp. concentration used during the phagocytic assay was inconclusive (Fig. 2A) presumably because algal cells might clumped together and the sponge became oversaturated during the incubation.

**Fig. 2.**
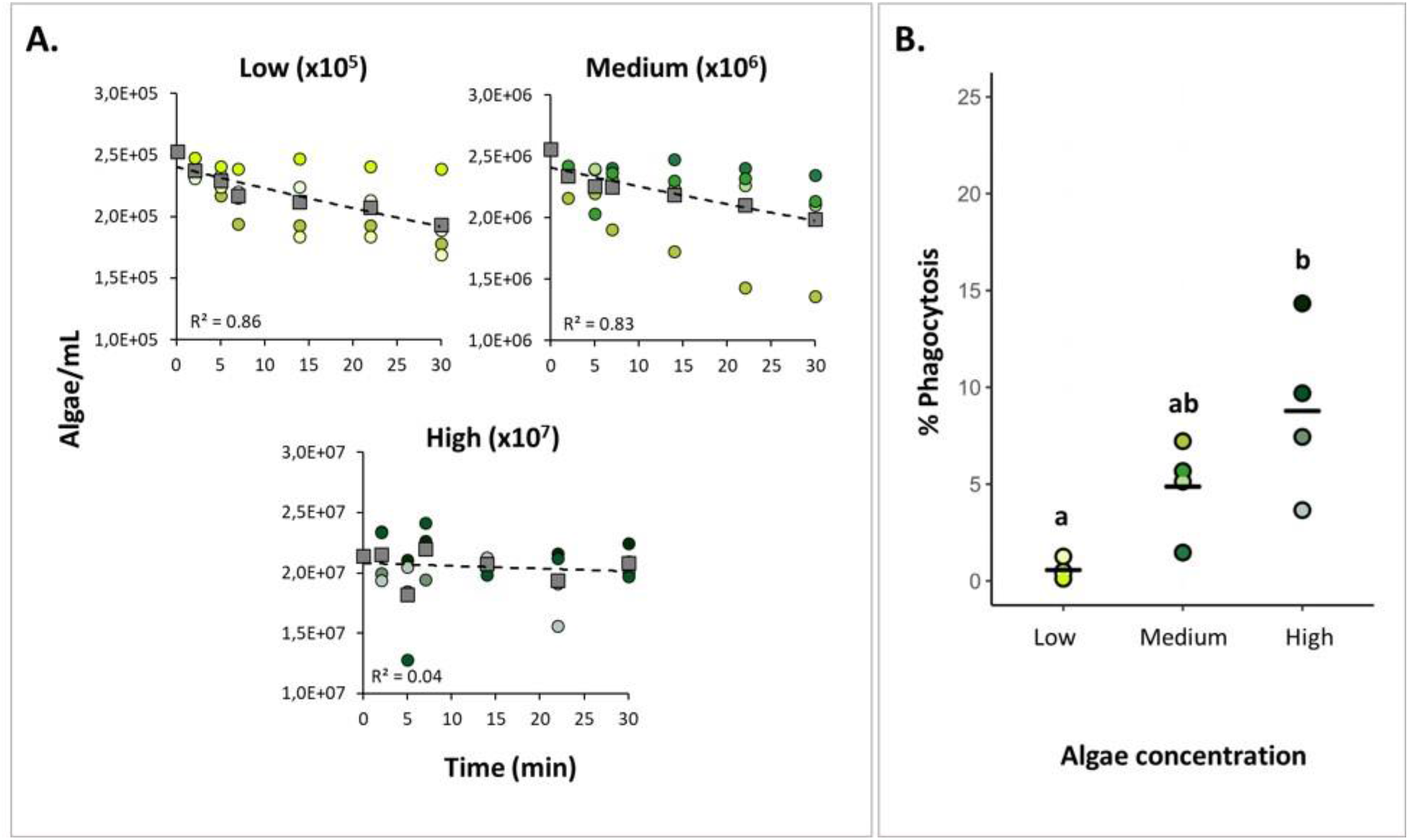
Testing tracer concentration for the phagocytic assay. **(A)** Algal uptake by *H. panicea* individuals incubated at three algal concentrations. Dots of the same color: biological replicates (n = 4 per treatment). Squares and dash lines: averaged data fitted to an exponential model. **(B)** Estimates of phagocytic active sponge cells. Bold line: average for the 4 biological replicates. Treatments marked with different letters are significantly different at α=0.05.

The percentage of phagocytic cells was 0.6 ± 0.4 % for the low concentration (×10^5^ cells mL^-1^), 4.9 ± 2.1 % for the medium concentration (×10^6^ cells mL^-1^), and 8.8 ± 3.9 for the high-concentration (×10^7^ cells mL^-1^) (Fig. 2B). There was a significant effect of algal concentration on the percentage of phagocytic cells in *H. panicea* (ANCOVA, F = 11.03, *p* < 0.01; df = 2). The medium and high algal concentrations yielded a 9-fold and 16-fold increase in the percentage of phagocytic cells compare to the lowest concentration, respectively. Whereas increasing the algal concentration from medium to high resulted in a 2-fold increase in the percentage of phagocytic cells (Fig. 2B). For the high concentration treatment, it is worth noting that even though the algal uptake estimates based on flow cytometry measurements were inconclusive, we did detect an increase in phagocytic activity, and observed incorporation of algae into the sponge cells.

### Assessing timing of algal phagocytosis

We performed a pulse-chase experiment using *Nannochloropsis* sp. to assess at what time point after presenting the algae to *H. panicea* yielded the highest percentage of phagocytic sponge cells. The total number of algal cells taken up by the sponges after the 30 min pulse period was 3.3 ± 2.5 × 10^5^ cells, which translates to a 13.7 ± 10.6 % algal reduction (Fig. 3A). The percentage of phagocytic cells was 4.9 ± 2.1 % after 0 min, 2.2 ± 0.78 % after 30 min, and 2.0 ± 0.4 % after 150 min chase-period (Fig. 3B). The percentage of phagocytic cells was positively related with algal uptake (ANCOVA, F = 6.53, *p* = 0.03; df = 1). After removing the effect of particle uptake on the response variable, phagocytic activity significantly decreased during the chase period (i.e., phagocytosis decreased with time after initial particle exposure), (ANCOVA, F = 9.80, *p* < 0.01; df = 2). The phagocytic activity peaked at 0 min chase, then significantly decreased by approx. 50 % at 30 min chase, but there was no further significant decline in the percentage of phagocytic cells after 150 min chase.

**Fig. 3.**
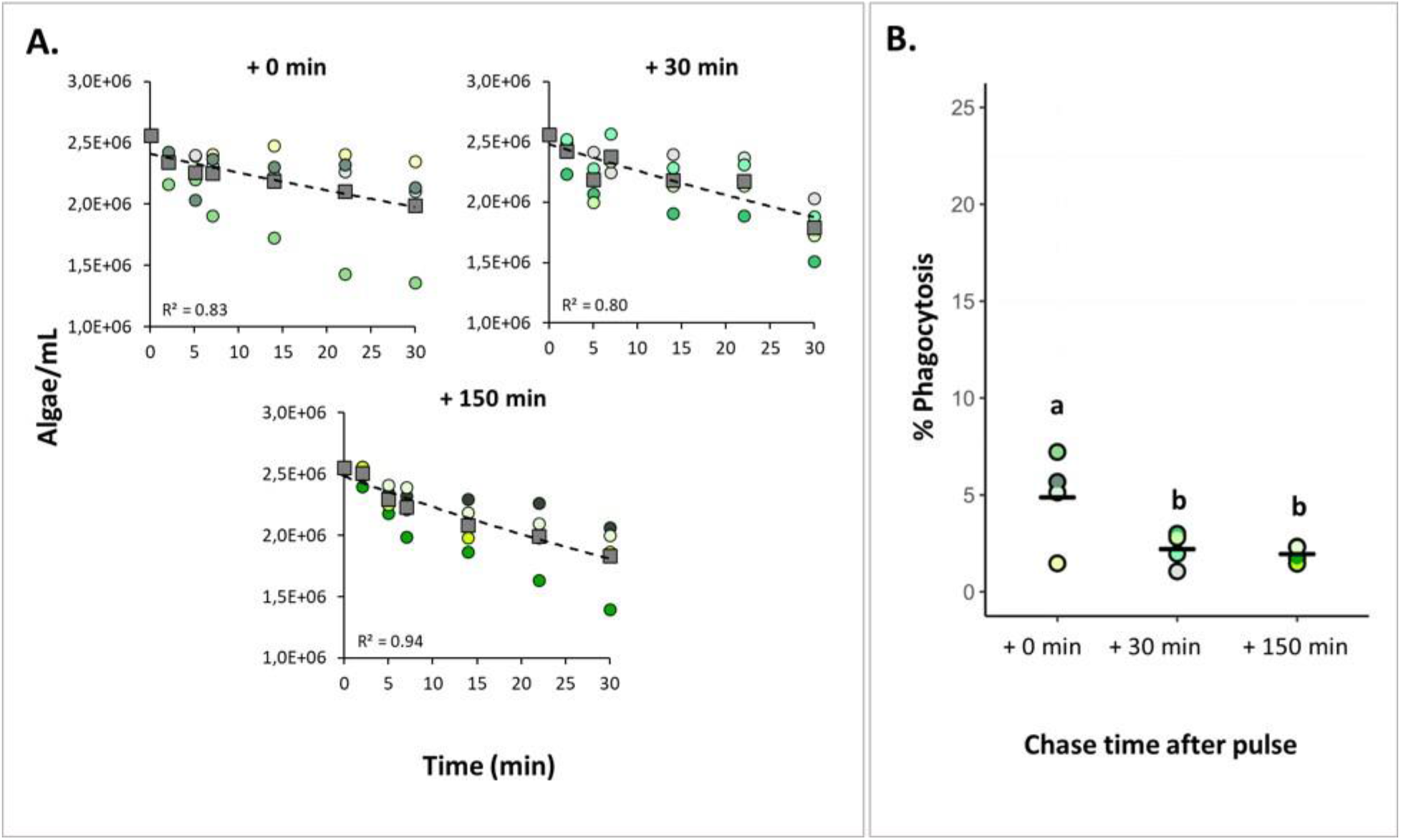
Assessing the starting time of algal phagocytosis. **(A)** Algal uptake by *H. panicea* individuals incubated for 30 min incubation (pulse period) and samples after three chase periods (+ 0 min, + 30 min, and + 150 min). Dots of the same color: biological replicates (n = 4 per treatment). Squares and dash lines: averaged data fitted to an exponential model. (**B)**. Estimates of phagocytic active sponge cells. Bold line: average for the 4 biological replicates. Treatments marked with different letters are significantly different at α=0.05.

### Bacteria and latex beads as tracers of phagocytosis

When bacteria and beads were provided as tracers, the total number of particles taken up by the sponge was 2.7 × 10^4^ ± 7.9 × 10^3^ bacteria and 2.4 × 10^5^ ± 2.3 × 10^5^ beads, corresponding to a 31.0 ± 8.6 % and 14.3 ± 16.0 % reduction after each assay, respectively (Fig. 4A). Particle uptake varied among *H. panicea* individuals. Some sponge individuals only reduced the particle concentration by 1 to 4 % during the assay, while in other assays the concentrations decreased by 22 to 37 %. Despite the observed variation, there was overall no significant difference in particle uptake between bacteria, beads, and algae (ANOVA, F = 0.57, *p* = 0.59; df = 2).

**Fig. 4.**
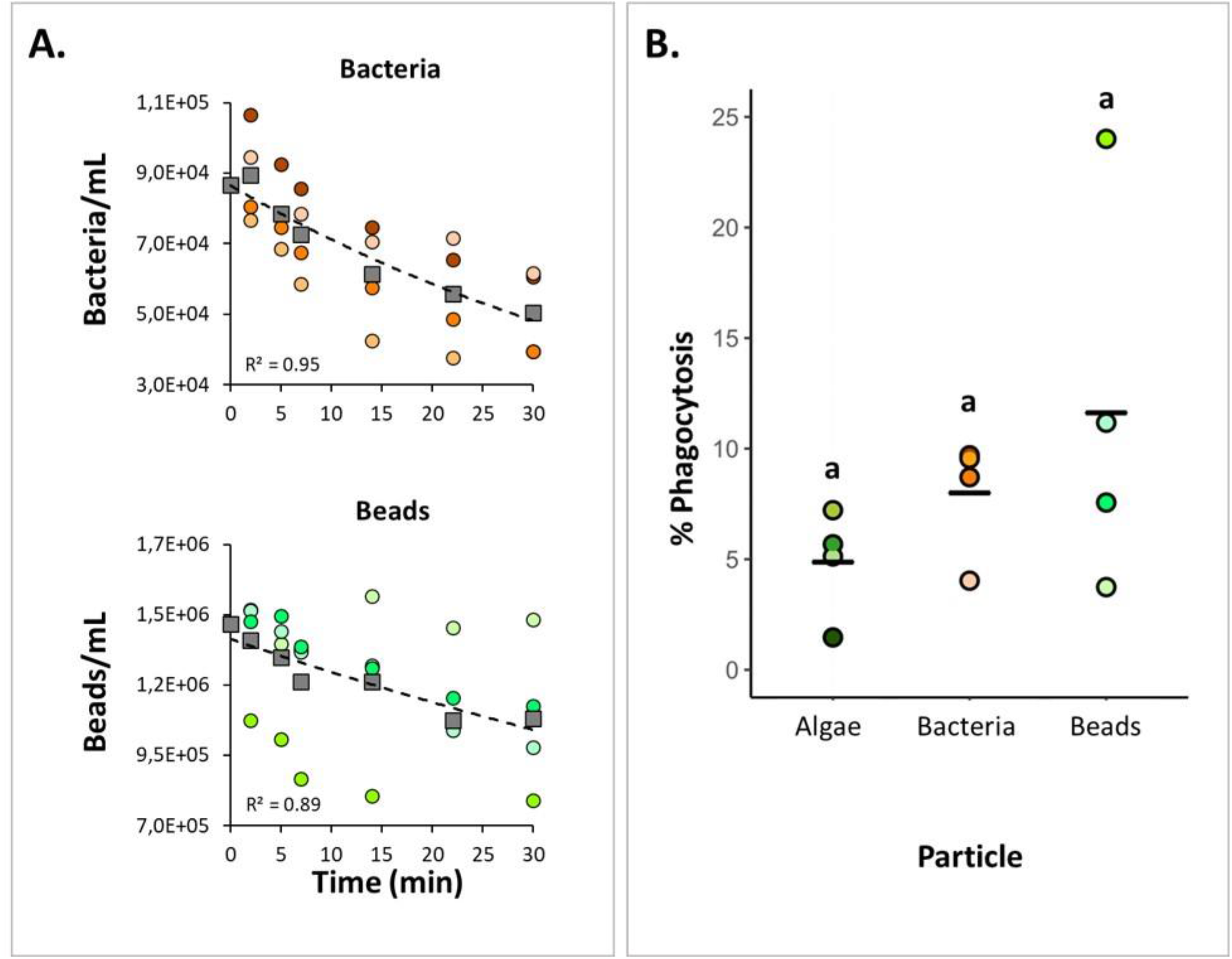
Testing bacteria and beads as tracers for the phagocytic assay. (**A)**. Bacterial and bead uptake by *H. panicea* individuals after 30 min incubation with TAMRA-stained bacteria (*Vibrio* sp.) and fluorescent latex beads (1 µm). Water samples for flow cytometry estimates were taken at time intervals. Dots of the same color: biological replicates (n = 4 per treatment). Squares and dash lines correspond to the averaged data fitted to an exponential model. **(B)** Estimates of phagocytic active sponge cells in comparison to the algal assay. Bold line: average for the 4 biological replicates. Treatments marked with different letters are significantly different at α=0.05.

In the bacteria experiment, sponge control samples (i.e., individuals incubated without bacteria) showed some events in the gate used for quantifying cells with incorporated fluorescent signals (Fig. 1C). Based on our microscopy observations using the filter set 43 HE DsRed (538-562 nm), the signal may correspond to natural auto-fluorescent granules present in the *H. panicea* cells. After subtracting the events estimated in the control samples, the percentage of cells phagocytizing bacteria in the treated sponges was 8.0 ± 2.3 % (Fig. 4B). In the assay with the beads, we identified a distinct population of phagocytic sponge cells consisting of at least three subpopulations (Fig. 1D). Fluorescent microscopy of the sorted cells within these three subpopulations revealed differences in the number of beads phagocytized per sponge cell. Most cells from the lower subpopulation in the y-axis (i.e., green fluorescence) showed incorporation of one bead, while cells from the medium and highest subpopulation contained 2-3 and > 3 beads per cell, respectively (Fig. 1D). Overall, the percentage of sponge cells phagocytizing beads was 11.6 ± 7.6 % (Fig. 4B). When comparing the phagocytic activity of *H. panicea* individuals exposed to bacteria, beads, and algae, we detected no significant difference between tracers after controlling for particle uptake (ANCOVA, F = 1.52, *p* = 0.28; df = 2).

### Cellular insights into the phagocytic process

Fluorescent microscopy of *H. panicea* dissociated cells revealed diversity in terms of morphology and size of the sponge cells engaged in phagocytosis (Fig. 5A-C). In general, we observed relatively small cells (approx. 5 µm) with a clear nucleus of around 2 µm and a flagellum of various length, which we presume are choanocytes. Medium to big cells (approx. 6 to 10 µm, and 10 to 12 µm, respectively) with a nucleus of around 5 µm, and no flagella that resemble archeocyte-like cells were also visible. For all particle types (i.e., algae, bacteria, and beads), 39 to 50 % of the cells performing phagocytosis were choanocytes (Table S1). However, medium to big phagocytic cells (10 to 12 µm) engaged more often in algal phagocytosis (29 and 18 % of total phagocytic cells, respectively) compared to bacteria (13 and 4 %, respectively) and beads (10 and 3 %, respectively) (Table S1).

**Fig. 5.**
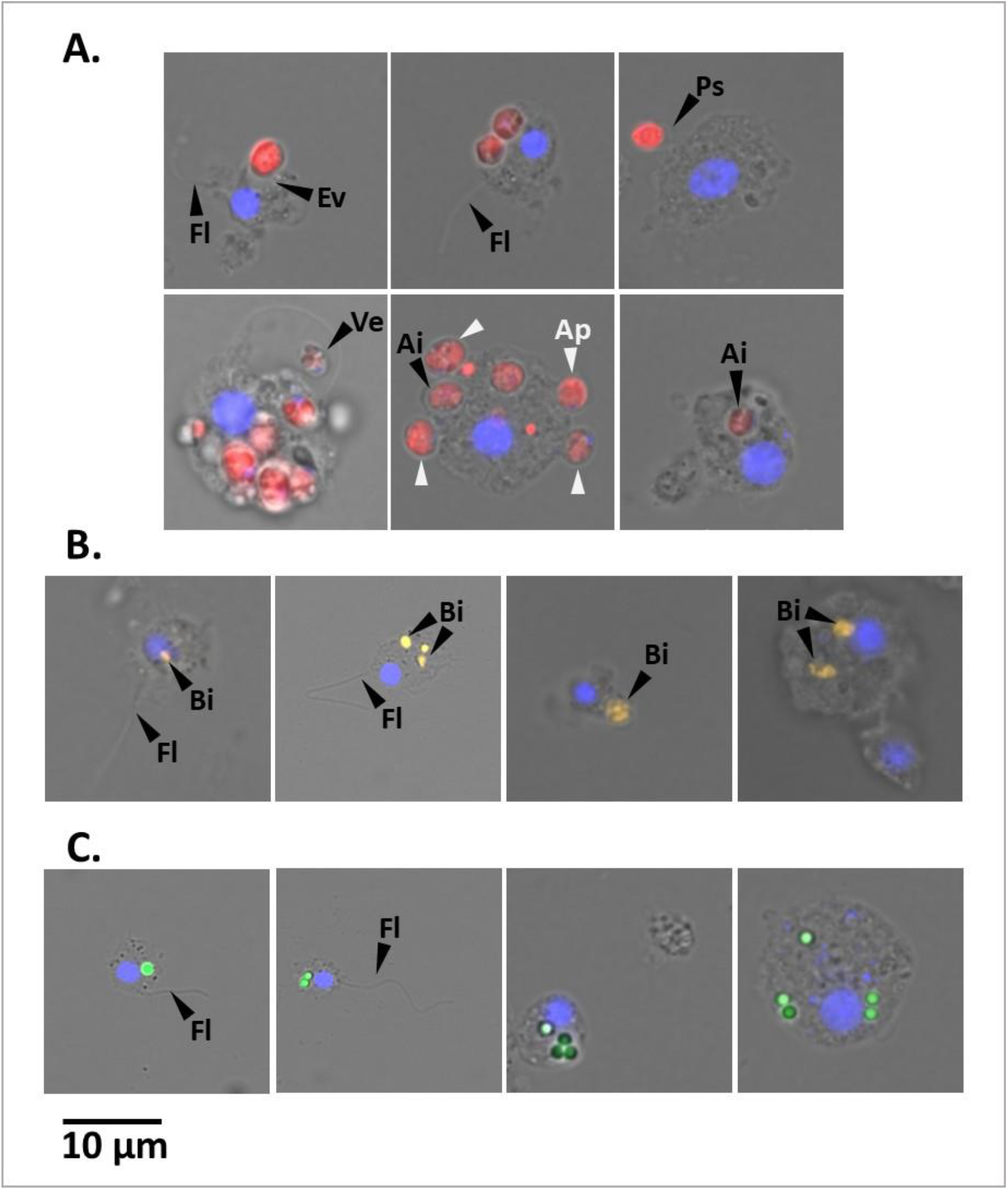
Representative fluorescent microscopy pictures of cells dissociated from *H. panicea* tissue after phagocytic assays using different tracers. Sponge cells phagocytizing **(A)** microalgae (*Nannochloropsis* sp.; red). Flagella (Fl); evagination (Ev) and pseudopodia (Ps) of sponge cell membrane for incorporating algae; algal cells in the periphery (Ap) and internalized (Ai) in the sponge cell; and vesicle (Ve) with algal cell. **(B)** TAMRA-stained *Vibrio* sp. (Vi; yellow) and **(C)** fluorescent latex beads (1 µm; green) internalized in different cell types. Sponge cell nuclei (blue) stained with DAPI. Scale bar: 10 µm in all cases.

In the pulse-chase experiment with alga incubations, we identified potential early stages of algal phagocytosis at + 0 min chase, in which the sponge cell membrane evaginates, protrudes into pseudopodia-like structures, or extends vesicles to incorporate *Nannochloropsis* sp. cells (Fig. 5A). Furthermore, the percentage of phagocytic choanocytes showed a 3 to 7-fold significant reduction (Table S1) one and three hours after the assay started (i.e., + 30 min and + 150 min time point, respectively). In contrast, the proportion of archeocyte-like cells continued to increase significantly by up to 30 % at + 150 min (Table S1).

TEM observations on *H. panicea* tissue samples from the algal assays provided additional evidence of *Nannochloropsis* sp. internalization and processing into the sponge cells. Intact algal cells were observed in the extracellular matrix of the sponge (i.e., the mesohyl), exhibiting characteristic structures like the cell wall, thylakoids, and vacuole (Fig. 6A). Whole *Nannochloropsis* sp. cells internalized in sponge cells were detected in the inspected samples (Fig. 6B-C). We also observed phagosome-like structures with potential remnants of cell wall and thylakoids, which we presume, is the result of algal digestion (Fig. 6D-F). Algal phagocytosis was distinctive from ‘natural’ bacterial phagocytosis (e.g., bacteria were found in smaller vacuoles; Fig. 6G-H) and was never observed in the inspected control sample.

**Fig. 6.**
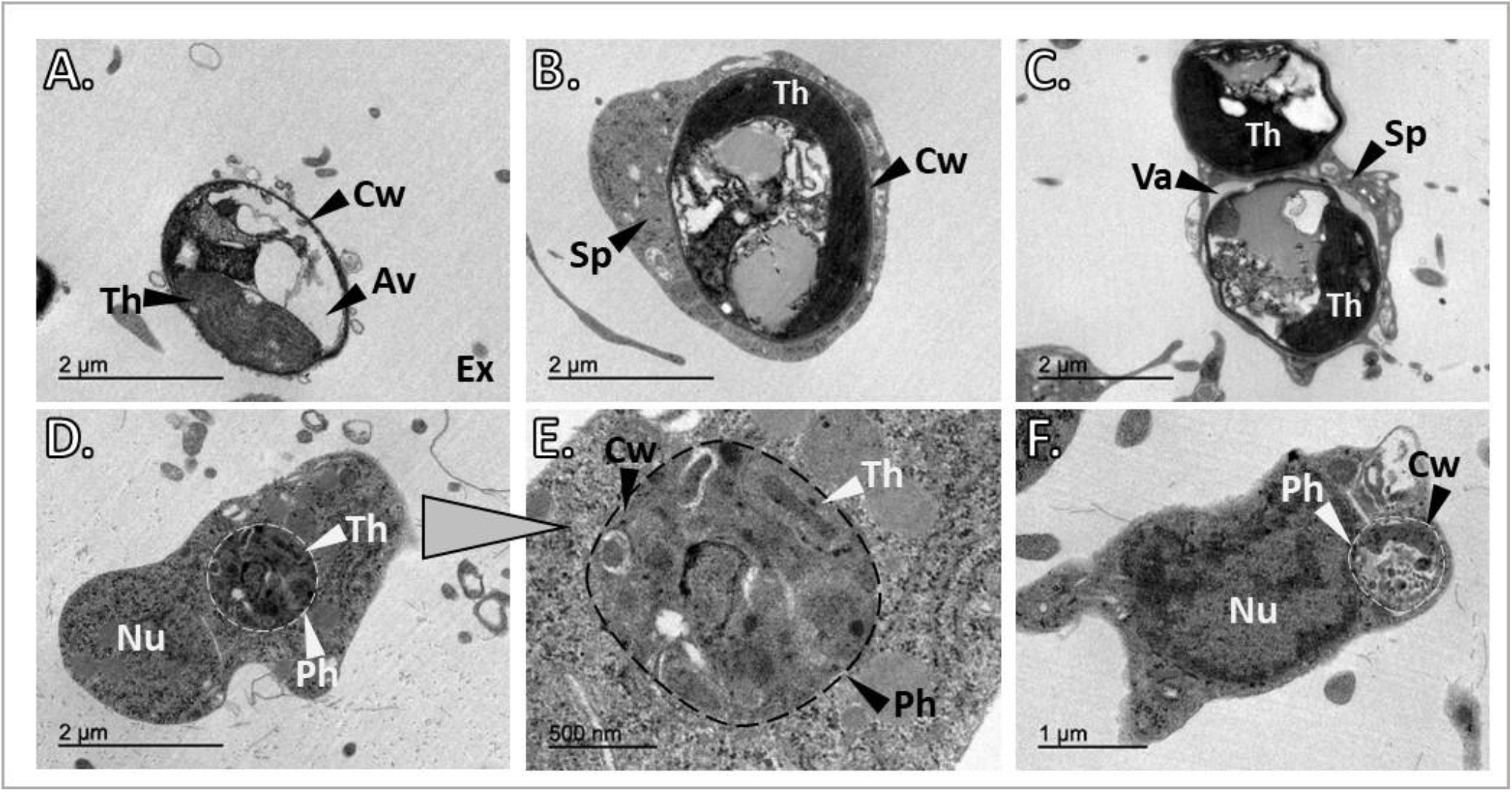
TEM observations from *H. panicea* tissue after algal phagocytosis. **(A)** Free, intact *Nannochloropsis* sp. cell in the sponge extracellular (Ex) matrix. The cell wall (Cw), thylakoids (Th), and the algae vacuole are visible. **(B)** and **(C)** sponge cells (Sp) with one and two internalized algal cells, respectively. One of the algae is inside a vacuole (Va). **(D)** to **(F)** Potential algal digestion in which possible remnants of algal cells are observed in phagosome-like structures (Ph). Nucleus (Nu).

## Discussion

In this study, we provide a novel approach to quantifying phagocytic active cells in whole sponge individuals by coupling live sponge incubations using fluorescent tracers, with sponge cell (host) dissociation, FACS analysis, and microscopy inspections. We adopted an in-vivo rather than an in-vitro cellular assay approach to replicate and accurately predict, as much as possible, the natural behavior of cells in *H. panicea*. In-vitro work in sponges is challenging mainly because their cells reaggregate fast, and the use of chemicals to induce disaggregation can reduce cell viability and inhibit essential cellular processes (46). With the established in-vivo phagocytic assay, we were able to successfully quantify the incorporation of algae, bacteria, and beads into *H. panicea* cells, and identify different sponge cells types involved in the process of phagocytosis. Overall, the developed phagocytic assay in *H. panicea* has enormous potential to be used as a tool for studying the role of phagocytosis for bacterial differentiation in sponge-microbe interactions.

### Tracer concentration and phagocytic activity

In each experiment, we quantified clearance of particles as indicator for sponge filtering activity (as in (47–50)). We observed that an increase in tracer concentration resulted in a linear increase in algal removal by *H. panicea*. This is consistent with previous findings of concentration-dependent particle removal rates in sponges (e.g., (15,48)). In our algal assays, most of the sponge individuals were actively filtering the tracer during the 30 min incubations with low and medium *Nannochloropsis* sp. concentrations (× 10^5^ and × 10^6^ cells mL^-1^, respectively), as the algal abundance declined exponentially over time. The algal uptake for the highest (× 10^7^ cells mL^-1^) *Nannochloropsis* sp. concentration used during the phagocytic assay was inconclusive. Algal cells might have clumped hindering the detection of changes in algal concentration during the assay and thus, we recommend working with particle concentrations below 10^7^ cells mL^-1^. The overall (mean ± SD) algal clearance rate of our sponges (1.5 ± 0.2 mL water mL sponge^-1^ min^-1^) is also in line with previous reports for *H. panicea* (1.5 to 1.9 ± 0.8 to 1.1 mL water mL sponge^-1^ min^-1^ (53)). Filtration rates of *H. panicea* can considerably vary in laboratory conditions (50), and even though we observed variation in the filtration activity of certain individuals, our incubation setup for the phagocytic assay proved overall to ensure the sponge filtration activity, which is an important factor to control.

We further hypothesized that the initial concentration of the tracer would also affect the estimates of phagocytic active cells. Indeed, we found a positive relation between algal concentration and the percentage of phagocytic cells. Increasing the algal concentration from 10^5^ to 10^6^ algae mL^-1^ resulted in a significant 8-fold increase in the sponge phagocytic activity. While the phagocytic activity seemed to further increase (2-fold) at an algal concentration of 10^7^ algae mL^-1^, this trend was not significant. This indicates that sponge cells can adjust their phagocytic activity depending on the number of algal cells they encounter. These results suggest that an algal concentration of approx. 10^6^ cells mL^-1^ is optimal for performing algal phagocytosis assays in *H. panicea*.

### Onset of phagocytosis

The observations from our pulse-chase experiment revealed internalization of algae into the sponge cells already 30 min (i.e., at + 0 min chase; Fig. 6B) after algal exposure, and after 1 h (i.e., + 30 min chase) phagocytic activity significantly decreased 2-fold, but then remained constant (i.e., + 150 min chase). In the chase phase, sponges were transferred to particle-free water, yet our FACS results showed that approx. 75 % of the algae that were taken up by the sponge were found as “intact algae” (i.e., present in the sponge fraction but not internalized in sponge cells) at + 0 min and + 30 min chase (Fig 6A and Fig S2). Our dissociation protocol was designed to be gentle enough to keep sponge cells intact and this was evident since we observed delicate structures like flagella (Fig 5). We suggest that intact algae are either algae that were loosely attached to sponge cells (e.g., Fig 5A) and got detached during the sponge cell dissociation process or during the FACS. Interestingly, at + 150 min chase the percentage of intact algae decreased to 56 % (Fig S2), supporting that intact *Nannochloropsis* sp. cells were taken up by the sponge during the pulse chase and were in the process of internalization during the chase phase.

We propose that the decrease in phagocytic activity between + 0 min chase and + 30 min chase is the consequence of algal digestion (e.g., Fig 6-TEM D-F), and during this process, incorporation of intact algae might decrease. Once digestion is completed the process of internalization could be resumed and hence, the maintenance of phagocytic activity between + 30 min and + 150 min chase. In freshwater sponges, algae internalization and translocation between sponge cells occurs within minutes after feeding, whereas digestion takes a couple of hours. For example, in young individuals of *Spongilla lacustris* hatched from gemmules, extensive digestion of algal cells by phagocytes was evident 5 h after feeding (4). Likewise, in algal-free (i.e., “aposymbiotic”) *Ephydatia muelleri* intracellular algal symbionts were observed to be internalized in archeocytes 4 h post-infection (51). Our findings together with the above observations reveal that algal phagocytosis in sponges initiates within minutes after particle exposure and takes a couple of hours to be completed. We further suggest that the fact that some *Nannochloropsis* sp. cells were taken up by *H. panicea* but not internalized, due potential digestion events of other algal cells, indicates a decoupling in time between algal uptake, internalization, and digestion by sponge cells.

### Algae, bacteria, and beads as tracers for the phagocytic assay

The percentage of phagocytic cells estimated with our assay for *H. panicea* using algae, bacteria, and beads as tracer particles ranged overall between 5 to 24 %. It is difficult to compare our estimates to other studies since, to our knowledge, similar quantifications of phagocytic sponge cells have not been done yet. However, approximate comparisons are possible against amoeba as they share similarities with sponge archeocytes and are well-established models for phagocytosis studies. For example, amoebal phagocytic activity of *Dictyostelium discodeium* upon exposure to GPF-tagged *Legionella pneumophila* was estimated to be < 2 % based on FACS counts (52), whereas in *Acanthamoeba casellanii* microscopy counts showed 15-35% of amoeba cells phagocytizing *E. coli* (25). In-vitro phagocytosis experiments in cnidarians, another closely phylogenetic-related group to sponges, estimated with FACS percentages of coral and sea anemone phagocytic cells between 2.2 to 7.8 % and 9 to 18 % after feeding cells with 1 µm latex beads and fluorescently labeled *Escherichia coli*, respectively (38,39). Although the aforementioned assays diverge from ours in the sense that they were performed in cell suspensions, under different conditions, and not in whole sponge individuals, the reported phagocytic activity in those studies is comparable to our estimates in *H. panicea*.

Our data suggest that phagocytosis differs depending on the sponge filtration activity (i.e., particle uptake) and on the type of particle used during the assay. The percentage of phagocytic cells tends to increase with increased particle uptake in all tracer types, but this relationship seems to be tracer-specific (Fig. S3). The increase in phagocytic cells was faster for beads, followed by bacteria, and at a lower degree for algae. The beads tend to accumulate in the cells (Fig. 1D and Fig. 5C) as they cannot be digested by the sponge. For *H. panicea*, we speculate that once the sponge cells are saturated with beads (≥ 5 beads per cell) new cells need to engage in the phagocytic process, which could explain the steeper increase in bead internalization. In *E. muelleri* highly accumulation of beads in the choanocytes occurs within 13 min of exposure to the particle, and after 15 min bead incorporation extends to the archeocytes (53). Thus, the fast uptake and high number of sponge cells with incorporated beads may be the result of the participation of more choanocytes within the first 30 min of exposure to the particles.

In the case of bacteria, the number of *Vibrio* cells incorporated per sponge cell was difficult to resolve because the resolution acquired with the fluorescent microscope was not high enough, and it was difficult to distinguish accurately our fluorescent bacteria from natural fluorescence occurring in the sponge cells. However, FACS allowed us to subtract the natural fluorescence based on the signal in control samples and we estimate that an approx. 50 % increase in bacterial uptake would yield a 50 % increase in the percentage of phagocytic cells (Fig. S3). NanoSIMS experiments in the marine encrusting sponge *Halisarca caerulea* using isotopically labeled bacteria indicate that bacteria are rapidly phagocytized by choanocytes (54). Within 15 min after feeding the sponge with labeled bacteria, 90 to 100 % of the choanocyte cells incorporated bacteria and individual bacteria cells were visible in intracellular vesicles. Moreover, bacterial digestion and assimilation processes can vary depending on the bacteria encountered by the sponge. For example, *Hymeniacidon perlevis* can rapidly process *E. coli*, whereas assimilation of *Vibrio anguillarum* is more laborious. *Vibrio* cells are semi-digested by choanocytes 4 h after feeding and digestion is further completed by amoebocytes (16). Our phagocytosis assay allows the quantification of bacterial incorporation into sponge cells and could be used to investigate the mechanistic bases of microbe recognition and differentiation by sponges. Combining the developed in-vivo assay with cell markers would aid to resolve which types of sponge cells activate a phagocytic response upon bacterial encounter, and whether the population of phagocytic cells change when the sponge is presented with different types of microorganisms.

In the case of the algae, the percentage of sponge cells engaged in phagocytosis was lower than in the other tracers. Algal phagocytosis only increased by 1.5 % despite an approx. 30 % increase in algal uptake (Fig. S3). It is plausible to suggest that the bigger size and cell wall structure of the *Nannochloropsis* cells requires more time for the sponge to digest the algae (see previous section for details). Overall, our results indicate that the phagocytic response of *H. panicea* depends on the nature of the particle the sponge encounters. Indigestible particles (i.e., latex beads) trigger a faster internalization into the sponge cells and likely involved different cell populations. Whereas for microorganism that can be digested (i.e., bacteria and algae), phagocytosis seems to vary with particle size. Big algal cells are internalized at a slower rate than smaller bacterial cells (Fig S3), and higher rate of translocation between sponge cell types is evident.

## Conclusion and perspectives

Here we present a novel, in-vivo assay to quantify phagocytic active sponge cells in whole *H. panacea* individuals. Coupling sponge incubations with sponge cell (host) dissociation and FACS analysis proved to be a suitable approach to track fluorescent tracers (i.e., algae, bacteria, and beads) from the surrounding water into the sponge cells, and to quantify the relative proportion of *H. panicea* cells engaged in phagocytosis. Furthermore, our method allowed us to identify characteristics of the phagocytosis process itself. *H. panicea* phagocytosis differs depending on the sponge filtration activity (i.e., particle uptake) and on the type of particle used during the assay. The number of particles incorporated and the degree of digestion by the sponge cells was tracer-dependent. Sponge phagocytosis is a fast process, initiating within minutes and concluding within < 60 min of exposure to the tracers, and involves different cell types based on our microscopy observations.

We envision our developed assay as a tool to query whether sponges process different microbes (e.g., food, symbiont, and pathogens) distinctively. In order to fully characterize the diversity of sponge cells capable of engaging in the phagocytic process developing of marker genes for marine sponges is needed. Fluorescent in situ hybridization probes (e.g., (6)) together with our experimental approach and FACS analysis could aid to identify sponge cell types responsible for different steps of the phagocytosis process, and to investigate if their activity changes after encountering different microbes or environmental stressors (e.g., elevated temperatures, ocean acidification, sedimentation) impair this process. Moreover, coupling single-cell RNA sequencing of populations of phagocytic cells could shed light on the molecular machinery behind this cellular process. Adapting our assay to other early evolutionary metazoans and model organisms (e.g., freshwater sponges, *Nemastostella* sp., and other cnidarians) could further help to better understand this conserved cellular mechanism and may therefore have the potential to unravel the role of phagocytosis in basal animal-microbe interactions.

## Author contributions

AMMG, LP and UH conceived the idea. AMMG planned and conducted the experiments. KB and AMMG performed the FACS analysis and fluorescent microscopy. LP performed the TEM inspections. The initial draft of the manuscript was written by AMMG and UH. All authors contributed to improving the article and approved the submitted version.

## Acknowledgments

We are grateful to Dr. Lara Schmittmann for the field collection support, and Dr. Ben Mueller for helpful discussions. We further acknowledge Sabrina Jung for technical assistance in the lab, Andrea Hethke for support with the cell dissociations, Fabian Wendt for technical support with the algae culture and experimental setup logistics, Janis Müller for his experimental support. We thank the TEM-SEM Electron Microscopy Unit from Scientific and Technological Centers (CCiTUB), Universitat de Barcelona, for their support and advice on TEM technique, and the International Max Planck Research School for Evolutionary Biology for supervision effort of AMMG.

## Supplementary material

The Supplementary Material for this article can be found online at: xxxx

## Data availability statement

The authors confirm that the data supporting the results and conclusions of this study are accessible withing the article. The complete estimates for particle uptake and phagocytic quantification can be found at https://www.pangaea.de (ticket submission: PDI-34177). If further data is required it would be available from the corresponding authors (LP and UH) on request.

## Supplementary figures

**Supplementary Figure 1.**
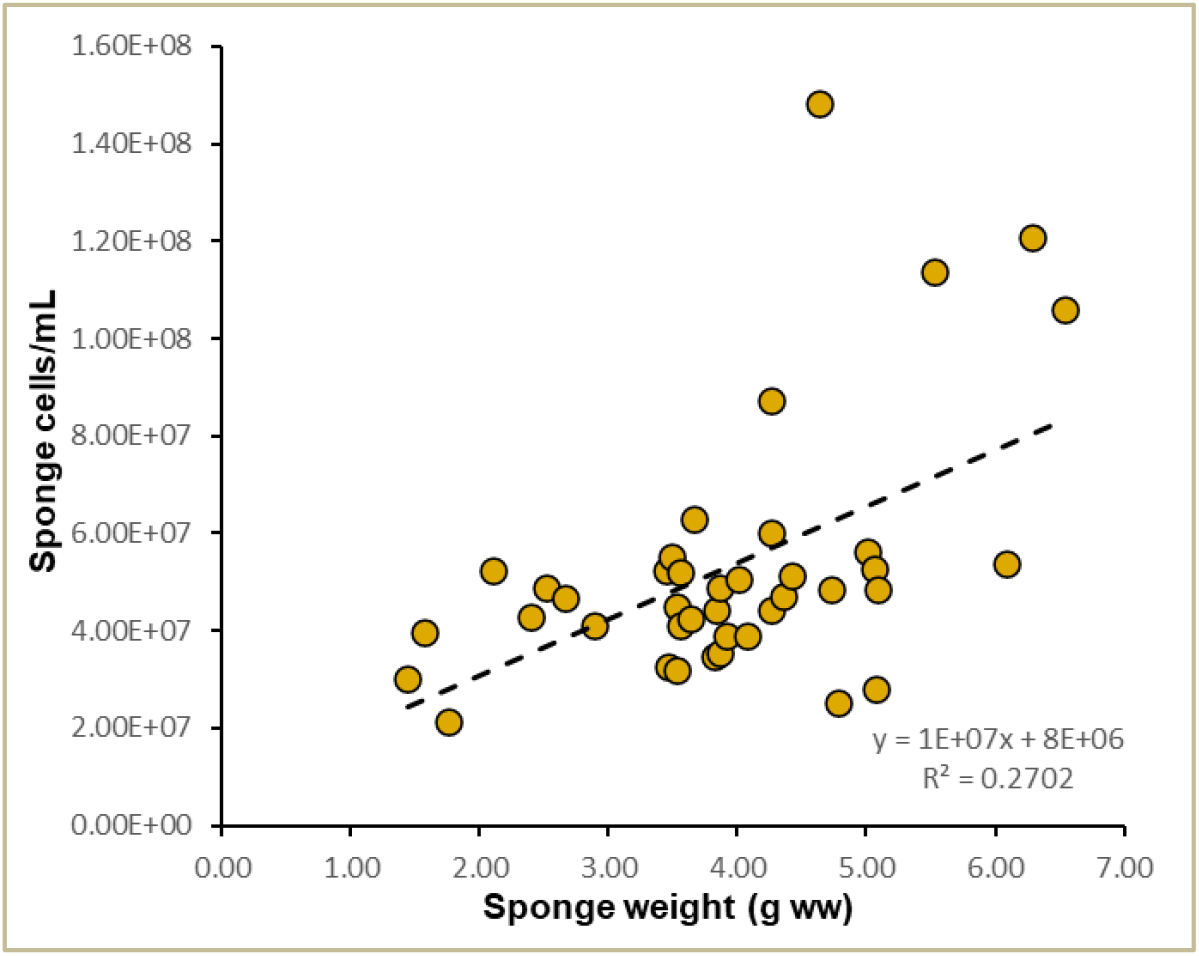
Relation of sponge cells recovered per g (wet weight) sponge after the cell dissociation of the individuals used for the phagocytosis assay.

**Supplementary Figure 2.**
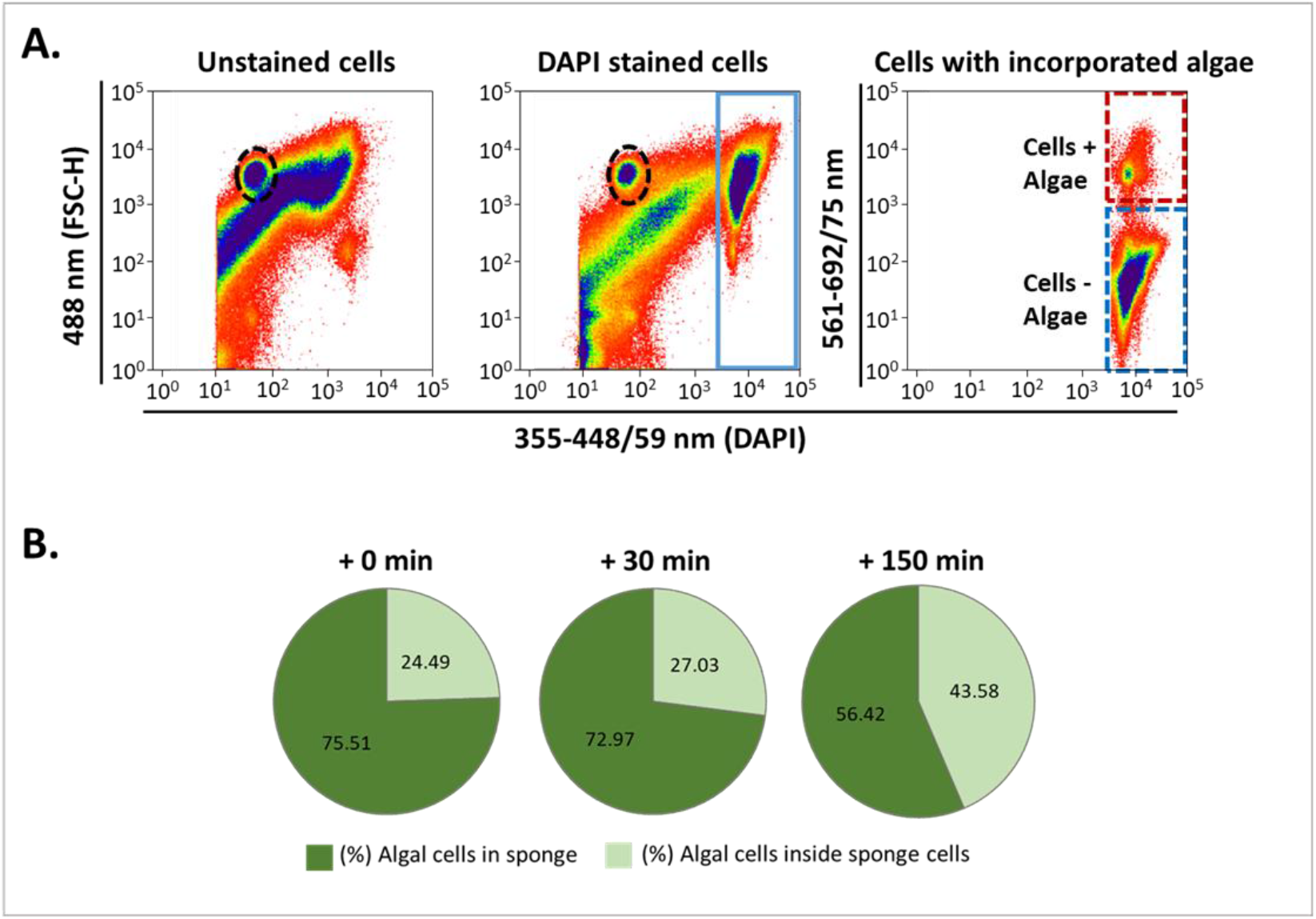
Estimates of *Nannochloropsis* sp. cells found inside the sponge tissue or incorporated in *H. panicea* cells **(A)** Representative FACS cytograms showing the population of intact, free algae (dashed oval). **(B)** Percentage of free algal cells after the pulse-chase experiment. All sponges were incubated for 30 min (pulse-period) with *Nannochloropsis* sp. and sampled after 0 min, 30 min, and 150 min chase period.

**Supplementary Figure 3.**
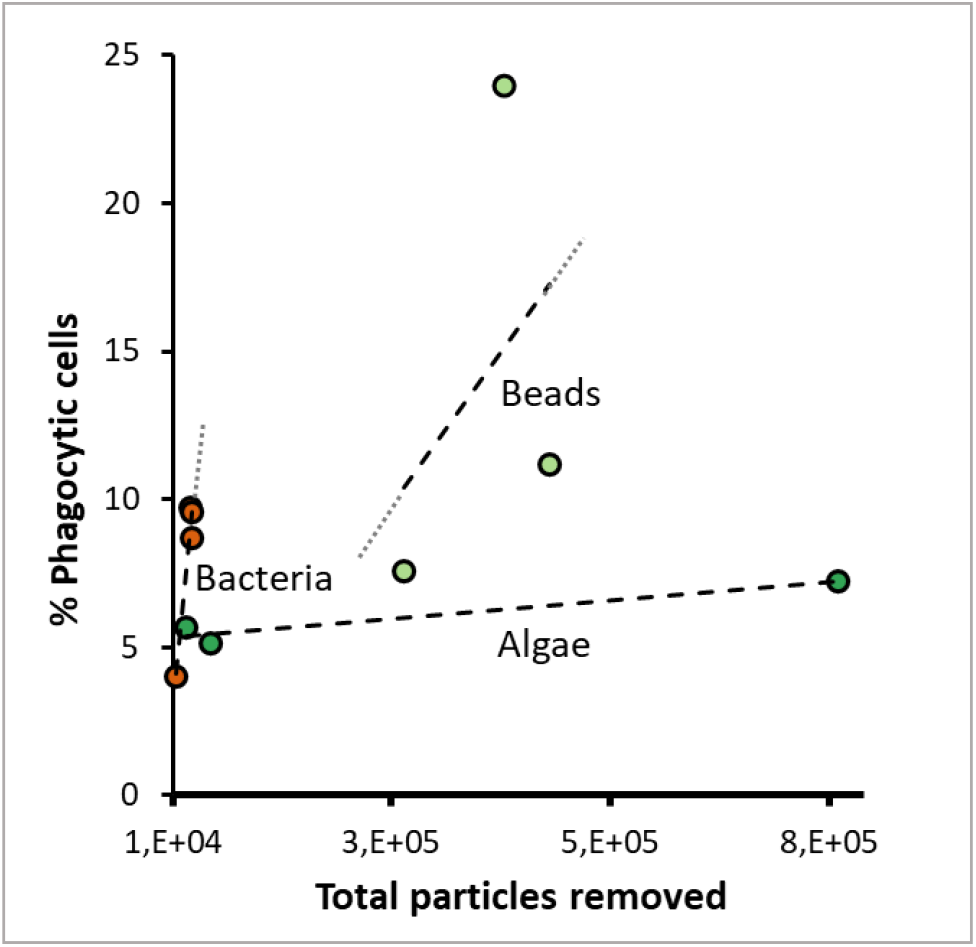
Relation between phagocytosis and particle uptake. The percentage of *H. panicea* cells phagocytizing algae (*Nannochloropsis* sp.), bacteria (*Vibrio* sp.), and fluorescent latex beads tends to increase linearly (n = 3. More data points are needed to validate a linear regression fit) with a higher number of particles removed by the sponge during the 30 min incubations. Dotted gray line: extension of the slope calculated with the linear regression equation.

**Supplementary Table 1.**
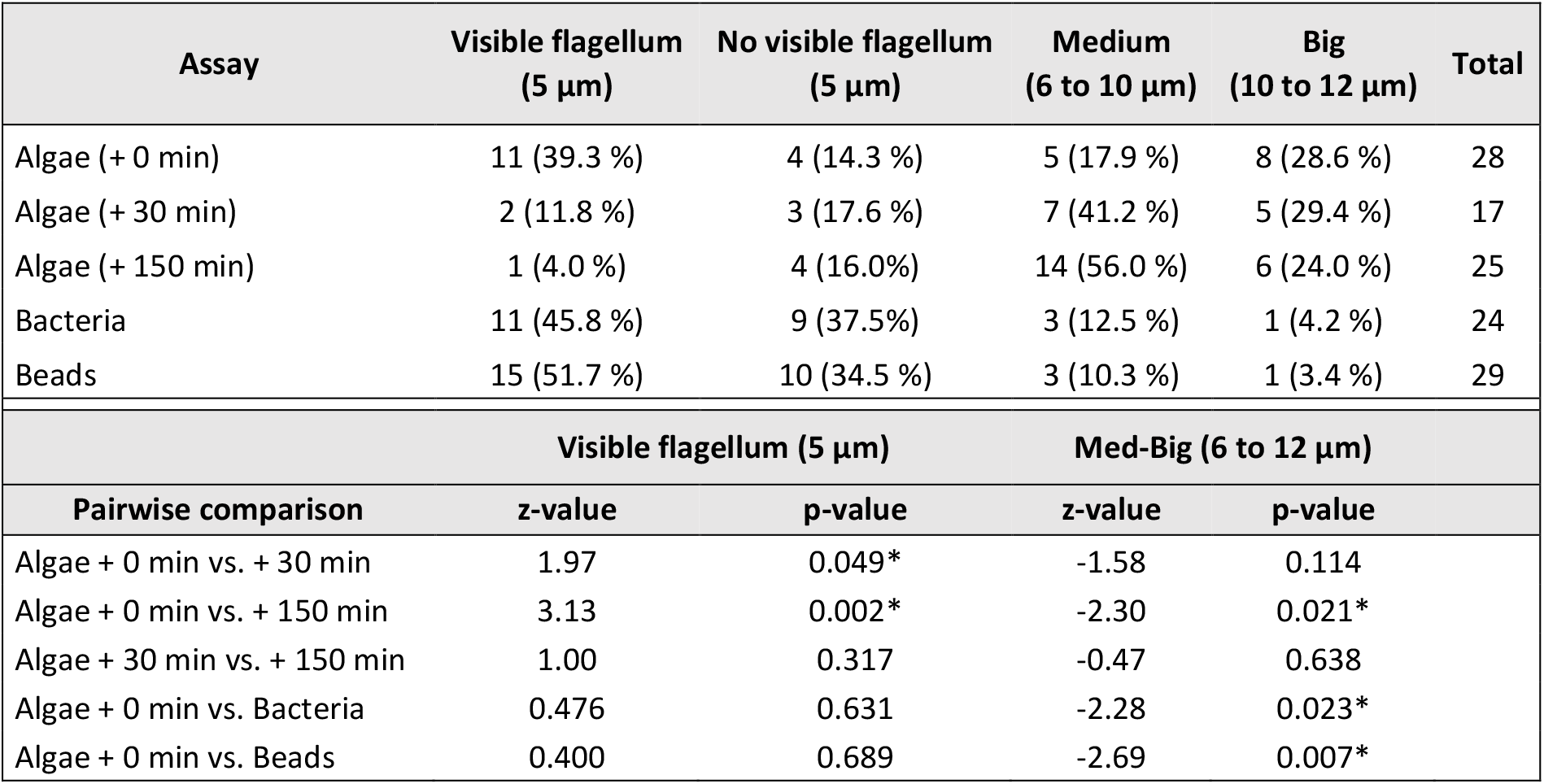
Fluorescent microscopy observations of phagocytic cells of *H. panicea* from the assays with algae (*Nannochloropsis* sp.), TAMRA-stained bacteria (*Vibrio* sp.), and fluorescent latex beads (1 µm). The total number of cells observed and percentages are reported. A two-tailed z-test was performed to compare the proportion of different cell types between treatments. *Significant values.

## References

1. Leys SP, Hill A. The physiology and molecular biology of sponge tissues. Adv Mar Biol (2012); 1–56. doi: 10.1016/B978-0-12-394283-8.00001-1

2. Simpson TL. The cell biology of sponges. Springer Verlag, New York (1984).

3. Steinmetz PR. A non-bilaterian perspective on the development and evolution of animal digestive systems. Cell Tissue Res (2019); 377(3):321–39. doi: 10.1007/s00441-019-03075-x

4. Imsiecke G. Ingestion, digestion, and egestion in Spongilla lacustris (Porifera, Spongillidae) after pulse feeding with Chlamydomonas reinhardtii (Volvocales). Zoomorphology (1993); 113(4):233–44. doi: 10.1007/BF00403314

5. Leys SP, Eerkes-medrano DI. Feeding in a calcareous sponge: particle uptake by pseudopodia. Biol Bull (2006); 211(2):157–71. doi: 10.2307/4134590.

6. Sebé-Pedrós A, Chomsky E, Pang K, Lara-Astiaso D, Gaiti F, Mukamel Z, Amit I, Hejnol A, Degnan BM, Tanay A. Early metazoan cell type diversity and the evolution of multicellular gene regulation. Nat Ecol Evol (2018); 2(7):1176–88. doi: 10.1038/s41559-018-0575-6

7. Musser JM, Schippers KJ, Nickel M, Mizzon G, Kohn AB, Pape C, Ronchi P, Papadopoulos N, Tarashansky AJ, Hammel JU, Wolf F. Profiling cellular diversity in sponges informs animal cell type and nervous system evolution. Science (2021); 374(6568):717–23. doi: 10.1126/science.abj294911.

8. Hentschel U, Usher KM, Taylor MW. Marine sponges as microbial fermenters. FEMS Microbiol Ecol. 2006;55(2):167–77.

9. Hentschel U, Usher KM, Taylor MW. Marine sponges as microbial fermenters. FEMS Microbio Ecol (2006); 55(2):167–77. doi: 10.1111/j.1574-6941.2005.00046.x

10. Thomas T, Moitinho-Silva L, Lurgi M, Björk JR, Easson C, Astudillo-García C, et al. Diversity, structure and convergent evolution of the global sponge microbiome. Nat Commun (2016); 7(1):11870. doi: 10.1038/ncomms11870

11. Moitinho-Silva L, Nielsen S, Amir A, Gonzalez A, Ackermann GL, Cerrano C, et al. The sponge microbiome project. Gigascience (2017); 6(10):1–7. doi: 10.1093/gigascience/gix077

12. Pile AJ, Young CM. The natural diet of a hexactinellid sponge: benthic-pelagic coupling in a deep-sea microbial food web. Deep Sea Res Part I Oceanogr Res Pap (2006); 53(7):1148–56. doi: 10.1016/j.dsr.2006.03.008

13. Reiswig HM. Particle feeding in natural populations of three marine demosponges. Biol Bull (1971);141(3):568–91.doi: 10.2307/1540270

14. Ribes M, Coma R, Gili JM. Natural diet and grazing rate of the temperate sponge Dysidea avara (Demospongiae, Dendroceratida) throughout an annual cycle. Mar Ecol Prog Ser (1999); 176:179–90.doi: 10.3354/meps176179

15. Turon X, Galera J, Uriz MJ. Clearance rates and aquiferous systems in two sponges with contrasting life-history strategies. J Exp Zool (1997); 278(1):22–36. doi: 10.1002/(SICI)1097-010X(19970501)278:1<22::AID-JEZ3>3.0.CO;2-8

16. Yahel G, Eerkes-Medrano DI, Leys SP. Size independent selective filtration of ultraplankton by hexactinellid glass sponges. Aquat Microb Ecol (2006); 24;45(2):181–94. doi:10.3354/ame045181

17. Maldonado M, Zhang X, Cao X, Xue L, Cao H, Zhang W. Selective feeding by sponges on pathogenic microbes: a reassessment of potential for abatement of microbial pollution. Mar Ecol Prog Ser (2010); 403:75-89. doi: 10.3354/meps08411

18. McMurray SE, Johnson ZI, Hunt DE, Pawlik JR, Finelli CM. Selective feeding by the giant barrel sponge enhances foraging efficiency. Limnol Oceanogr (2016); 61(4):1271–86. doi: 10.1002/lno.10287

19. Wilkinson CR, Garrone R, Vacelet J. Marine sponges discriminate between food bacteria and bacterial symbionts: electron microscope radioautography and in situ evidence. Proc R Soc London - Biol Sci (1984); 220(1221):519–28. doi: 10.1098/rspb.1984.0018

20. Wehrl M, Steinert M, Hentschel U. Bacterial uptake by the marine sponge Aplysina aerophoba. Microb Ecol (2007); 53(2):355–65. doi: 10.1007/s00248-006-9090-4

21. Nyholm S V., Graf J. Knowing your friends: invertebrate innate immunity fosters beneficial bacterial symbioses. Nat Rev Microbiol (2012) ;10(12):815–27. doi: 10.1038/nrmicro2894

22. Rosales C, Uribe-Querol E. Phagocytosis: a fundamental process in immunity. Biomed Res Int (2017); 2017. doi: 10.1155/2017/9042851

23. Hartenstein V, Martinez P. Phagocytosis in cellular defense and nutrition: a food-centered approach to the evolution of macrophages. Cell Tissue Res (2019); 377(3):527–47. doi: 10.1007/s00441-019-03096-6

24. Lim JJ, Grinstein S, Roth Z. Diversity and versatility of phagocytosis: roles in innate immunity, tissue remodeling, and homeostasis. Front Cell Infect Microbiol (2017); 7:1–12. doi: 10.3389/fcimb.2017.00191

25. Jahn MT, Arkhipova K, Markert SM, Stigloher C, Lachnit T, Pita L, et al. A phage protein aids bacterial symbionts in eukaryote immune evasion. Cell Host Microbe (2019); 26(4):542–550. doi: 10.1016/j.chom.2019.08.019

26. Nguyen Mthd, Liu M, Thomas T. Ankyrin-repeat proteins from sponge symbionts modulate amoebal phagocytosis. Mol Ecol (2014); 23(6):1635–45. doi: 10.1111/mec.12384

27. Nyholm S V., Stewart JJ, Ruby EG, McFall-Ngai MJ. Recognition between symbiotic Vibrio fischeri and the haemocytes of Euprymna scolopes. Environ Microbiol (2009); 11(2):483–93. doi: 10.1111/j.1462-2920.2008.01788.x

28. Silver AC, Kikuchi Y, Fadl AA, Sha J, Chopra AK, Graf J. Interaction between innate immune cells and a bacterial type III secretion system in mutualistic and pathogenic associations. Proc Natl Acad Sci (2007); 104(22):9481–6. doi: 10.1073/pnas.0700286104

29. Thomas T, Rusch D, DeMaere MZ, Yung PY, Lewis M, Halpern A, et al. Functional genomic signatures of sponge bacteria reveal unique and shared features of symbiosis. ISME J (2010); 4(12):1557–67. doi: 10.1038/ismej.2010.74

30. Díez-Vives C, Moitinho-Silva L, Nielsen S, Reynolds D, Thomas T. Expression of eukaryotic-like protein in the microbiome of sponges. Mol Ecol (2017); 26(5):1432–51. doi: 10.1111/mec.14003

31. Siegl A, Kamke J, Hochmuth T, Piel J, Richter M, Liang C, et al. Single-cell genomics reveals the lifestyle of Poribacteria, a candidate phylum symbiotically associated with marine sponges. ISME J (2011); 5(1):61–70. doi: 10.1038/ismej.2010.95

32. Al-Khodor S, Price CT, Habyarimana F, Kalia A, Abu Kwaik Y. A Dot/Icm-translocated ankyrin protein of Legionella pneumophila is required for intracellular proliferation within human macrophages and protozoa. Mol Microbiol (2008); 70(4):908–23. doi: 10.1111/j.1365-2958.2008.06453.x

33. Habyarimana F, Al-Khodor S, Kalia A, Graham JE, Price CT, Garcia MT, et al. Role for the Ankyrin eukaryotic-like genes of Legionella pneumophila in parasitism of protozoan hosts and human macrophages. Environ Microbiol (2008); 10(6):1460–74. doi: 10.1111/j.1462-2920.2007.01560.x

34. Pan X, Lührmann A, Satoh A, Laskowski-Arce MA, Roy CR. Ankyrin repeat proteins comprise a diverse family of bacterial type IV effectors. Science (2008); 320(5883):1651–4. doi: 10.1126/science.1158160

35. Sattler N, Monroy R, Soldati T. Quantitative analysis of phagocytosis and phagosome maturation. Methods Mol Biol (2013); 983:383–402. doi: 10.1007/978-1-62703-302-2_21

36. Jauslin T, Lamrabet O, Crespo-Yañez X, Marchetti A, Ayadi I, Ifrid E, et al. How phagocytic cells kill different bacteria: a quantitative analysis using Dictyostelium discoideum. MBio (2021); 12(1):1–13. doi: 10.1128/mBio.03169-20

37. Bucher M, Wolfowicz I, Voss PA, Hambleton EA, Guse A. Development and symbiosis establishment in the Cnidarian endosymbiosis model Aiptasia sp. Sci Rep (2016); 6:1–11. doi: 10.1038/srep19867

38. Jacobovitz MR, Rupp S, Voss PA, Maegele I, Gornik SG, Guse A. Dinoflagellate symbionts escape vomocytosis by host cell immune suppression. Nat Microbiol (2021); 6(6):769–82. doi: 10.1038/s41564-021-00897-w

39. Rosental B, Kozhekbaeva Z, Fernhoff N, Tsai JM, Traylor-Knowles N. Coral cell separation and isolation by fluorescence-activated cell sorting (FACS). BMC Cell Biol (2017); 18(1):1–12. doi: 10.1186/s12860-017-0146-8

40. Snyder GA, Eliachar S, Connelly MT, Talice S, Hadad U, Gershoni-yahalom O, et al. Functional characterization of Hexacorallia phagocytic cells. Front Immunol (2021); 12:1–13. doi: 10.3389/fimmu.2021.662803

41. Gasol JM, Del Giorgio PA. Using flow cytometry for counting natural planktonic bacteria and understanding the structure of planktonic bacterial communities. Sci Mar (2000); 64(2):197–224. doi: 10.3989/scimar.2000.64n2197

42. Fieseler L, Horn M, Wagner M, Hentschel U. Discovery of the novel candidate phylum “Poribacteria” in marine sponges. Appl Environ Microbiol (2004); 70(6):3724–32. doi: 10.1128/AEM.70.6.3724-3732.2004

43. Scheffers SR, Nieuwland G, Bak RPM, Van Duyl FC. Removal of bacteria and nutrient dynamics within the coral reef framework of Curaçao (Netherlands Antilles). Coral Reefs (2004); 23(3):413–22. doi: 10.1007/s00338-004-0400-3

44. de Goeij JM, Van Duyl FC. Coral cavities are sinks of dissolved organic carbon (DOC). Limnol Oceanogr (2007); 52(6):2608–17. doi: 10.4319/lo.2007.52.6.2608

45. Yahel G, Whitney F, Reiswig HM, Eerkes-Medrano DI, Leys SP. In situ feeding and metabolism of glass sponges (Hexactinellida, Porifera) studied in a deep temperate fjord with a remotely operated submersible. Limnol Oceanogr (2007); 52(1):428–40. doi: 10.4319/lo.2007.52.1.0428

46. de Goeij JM, van Den Berg H, van Oostveen MM, Epping EHG, Van Duyl FC. Major bulk dissolved organic carbon (DOC) removal by encrusting coral reef cavity sponges. Mar Ecol Prog Ser (2008); 357:139–51. doi: 10.3354/meps07403

47. Pomponi SA. Biology of the Porifera: cell culture. Vol. 84, Can J Zool (2006); 84(2):167–74. doi: 10.1139/z05-188

48. Leys SP, Yahel G, Reidenbach MA, Tunnicliffe V, Shavit U, Reiswig HM. The sponge pump: the role of current induced flow in the design of the sponge body plan. PLoS One (2011); 6(12). doi: 10.1371/journal.pone.0027787

49. Mueller B, de Goeij JM, Vermeij MJA, Mulders Y, Van Der Ent E, Ribes M, et al. Natural diet of coral-excavating sponges consists mainly of dissolved organic carbon (DOC). PLoS One (2014); 9(2). doi: 10.1371/journal.pone.0090152

50. Lüskow F, Riisgård HU, Solovyeva V, Brewer JR, Kløve-Mogensen K, Tophøj J, et al. Seasonal changes in bacteria and phytoplankton biomass control the condition index of the demosponge Halichondria panicea in temperate Danish waters. Mar Ecol Prog Ser (2019); 6:1–7. doi: 10.3354/meps12785

51. Riisgård HU, Kumala L, Charitonidou K. Using the F/R-ratio for an evaluation of the ability of the demosponge Halichondria panicea to nourish solely on phytoplankton versus free-living bacteria in the sea. Mar Biol Res (2016); 12(9):907–16. doi: 10.1080/17451000.2016.1206941

52. Geraghty S, Koutsouveli V, Hall C, Chang L, Sacristan-Soriano O, Hill M, et al. Establishment of host-algal endosymbioses: genetic response to symbiont versus prey in a sponge host. Genome Biol Evol (2021); 13(11):1–16. doi: 10.1093/gbe/evab252

53. Hägele S, Köhler R, Merkert H, Schleicher M, Hacker J, Steinert M. Dictyostelium discoideum: a new host model system for intracellular pathogens of the genus Legionella. Cell Microbiol (2000); 2(2):165–71. doi: 10.1046/j.1462-5822.2000.00044.x

54. Funayama N, Nakatsukasa M, Hayashi T, Agata K. Isolation of the choanocyte in the fresh water sponge, Ephydatia fluviatilis and its lineage marker, Ef annexin. Dev Growth Differ (2005); 47(4):243–53. doi: 10.1111/j.1440-169X.2005.00800.x

55. Hudspith M, Rix L, Achlatis M, Bougoure J, Guagliardo P, Clode PL, et al. Subcellular view of host–microbiome nutrient exchange in sponges: insights into the ecological success of an early metazoan–microbe symbiosis. Microbiome (2021); 9(1):1–15. doi: 10.1186/s40168-020-00984-w

